# Relating quantitative 7T MRI across cortical depths to cytoarchitectonics, gene expression and connectomics: a framework for tracking neurodegenerative disease

**DOI:** 10.1101/2020.02.05.935080

**Authors:** Peter McColgan, Saskia Helbling, Lenka Vaculčiaková, Kerrin Pine, Konrad Wagstyl, Fakhereh Movahedian Attar, Luke Edwards, Marina Papoutsi, Yongbin Wei, Martijn Pieter Van den Heuvel, Sarah Tabrizi, Geraint Rees, Nikolaus Weiskopf

## Abstract

Cortical layer-specific ultra-high field MRI has the potential to provide anatomically precise biomarkers and mechanistic insights into neurodegenerative disease. Here we compare cortical layer-specificity for a 7T multi-parametric mapping (MPM) 500μm whole brain acquisition to the von Economo and Big Brain post-mortem histology atlases. We also investigate the relationship between 7T MPMs, layer-specific gene expression and Huntington’s disease related genes, using the Allen Human Brain atlas. Finally we link MPM cortical depth measures with white matter connections using high-fidelity diffusion tractography from a 300mT/m Connectom MRI system. We show that R2* across cortical depths is highly correlated with layer-specific cell number, cell staining intensity and gene expression. Furthermore white matter connections were highly correlated with grey matter R1 and R2* across cortical depths. These findings demonstrate the potential of combining 7T MPMs, gene expression and white matter connections to provide an anatomically precise framework for tracking neurodegenerative disease.

## Introduction

The advent of ultra high field (UHF) MRI now enables us to image the human brain at sub-millimetre resolution in-vivo. Combining this technological advance with quantitative MRI (qMRI) has now made in-vivo histology MRI (hMRI) a distinct possibility (Trampel et al., 2019). Multi-parametric maps (MPMs) include qMRI parameters of effective transverse relaxation rate (R2*), which are sensitive to both myelin and iron (Weiskopf et al., 2013; Edwards et al., 2018) and longitudinal relaxation rate (R1), which is mainly sensitive to myelin and to a lesser extent to iron (Stuber et al., 2014).

The human cerebral cortex is composed of distinct cytoarchitectonic cortical layers, which are defined based on cell density, cell size and cell type. In-vivo high-resolution histology using UHF qMRI has the potential to provide cortical layer-specific measures that directly relate to these patterns of cell composition and associated layer-specific gene expression. Achieving in-vivo layer-specificity would further our understanding of the relationship between brain microstructure and function and how this is associated with sensory, motor and cognitive processing. In-vivo high-resolution UHF qMRI could also enable us to investigate neurodegenerative disease with much greater anatomical precision. For example in Alzheimer’s disease, superficial cortical layers show greatest vulnerability (Romito-DiGiacomo et al., 2007; Busche et al., 2008). However in both Parkinson’s disease (Pasquereau et al., 2016) and amyotrophic lateral sclerosis (Braak et al., 2017) deep cortical layers are selectively vulnerable, while in end stage Huntington’s disease post-mortem studies show involvement of layers 3, 5 and 6 (Rub et al., 2016). Thus layer-specific in-vivo MRI has the potential to provide mechanistic insights into layer selective vulnerability, as well as anatomically precise cortical biomarkers. In this study, we aim to link and characterize the sensitivity of layer-specific cortical measures using 7T qMRI in healthy humans in-vivo to established histological and gene expression measures. For this purpose, we relate R1 and R2* at different cortical depths from high-resolution MPMs to post-mortem whole brain histology atlases, gene enrichment atlases and whole brain connectomics derived from diffusion MRI. This is an important step in the development of UHF qMRI as a tool for in-vivo histology.

Here we focus on two post-mortem whole brain histology atlases that provide layer-specific quantitative cell measures, the von Economo Koskinas atlas and the Big Brain atlas. Von Economo and Koskinas published their seminal work the “Atlas of Cyto-architectonics of the Adult Human Cerebral Cortex” in 1925. They parcellated the cerebral cortex into 56 regions based on cell type, cell size and cell count (von Economo, 1925). By translating the von Economo regions into comparable Freesurfer Desikan regions (Scholtens et al., 2015) demonstrated highly significant correlation with post-mortem cortical thickness and in-vivo MRI cortical thickness. Subsequently the von Economo regions themselves have been mapped to MRI template space (Scholtens et al., 2018; van den Heuvel et al., 2019). This enabled us to directly test the sensitivity of UHF qMRI parameters to cytoarchitecture.

The Big Brain atlas (Amunts et al., 2013) was created more recently. The brain of a 65 year-old neurotypical male was sectioned into 20μm slices, stained for cell bodies and reconstructed in 3-D. Machine learning approaches (Wagstyl et al., 2018; Wagstyl et al., 2019) have been employed to define cortical layers in the Big Brain data providing a comparable cortical layer histology atlas to the von Economo atlas. Using the von Economo and Big Brain atlases we could therefore test the hypothesis that UHF qMRI parameters near the pial surface are correlated with cell measures in superficial layers 1-3, UHF qMRI parameters at mid-cortical depth are correlated with layer 4 and UHF qMRI parameters near the grey matter/white matter (GM/WM) boundary are correlated with deep cortical layers 5-6.

Going beyond cell histology, the Allen Human Brain Atlas (AHBA) provides a densely sampled atlas of regional gene expression across the human brain (Hawrylycz et al., 2012). For the AHBAsix brains underwent post-mortem MRI scanning enabling the mapping of gene expression to in-vivo MRI data. This allowed us to link qMRI parameters across cortical depths with layer-specific gene expression (Burt et al., 2018) and genes associated with disease pathophysiology (Rittman et al., 2016; McColgan et al., 2018). We could then test the hypothesis that UHF qMRI parameters across cortical depths are related to layer-specific gene expression in keeping with our hypothesis for UHF qMRI parameters and cytoarchitecture. We also investigated the relationship between regional UHF qMRI parameters and the regional expression of genes implicated in Huntington’s disease (HD) pathogenesis as an indicator of the potential application of these measures in neurodegeneration.

To demonstrate the sensitivity of UHF qMRI parameters to anatomical variability across cortical depths we related them to white matter connections. Based on *a priori* anatomical knowledge intratelencephalic neurons are found in cortical layers 2-6 and project to other cortical regions and to the striatum, forming cortico-cortical and cortico-striatal white matter connections, while pyramidal tract neurons are found in layer 5 and project to the striatum, forming cortico-striatal connections and also connections to deeper structures. Cortico-thalamic neurons are found in layer 6 forming cortico-thalamic connections (Molyneaux et al., 2007; Shepherd, 2013). We therefore hypothesised that cortico-cortical connections would be highly correlated with UHF qMRI parameters across all cortical depths, while cortico-striatal and cortico-thalamic connections would be highly correlated with UHF qMRI parameters near the grey matter/white matter (GM/WM) boundary.

In order to test these hypotheses we acquired whole brain MPMs at 7T with 500μm resolution in a group of 10 healthy young adults. We first present the R1 and R2* profiles across cortical depths. We then investigate the relationship of these measures across cortical depths with von Economo cortical layer cell count, Big Brain cell staining intensity and layer-specific gene expression using the AHBA. In addition we use the AHBA to explore the relationship of these whole brain qMRI parameters with expression of genes implicated in HD. Finally, we link cortical depth UHF qMRI parameters with specific anatomical *a priori*-defined white matter connections using diffusion tractography and connectomics, providing a high anatomical precision framework that can be used in future to track neurodegenerative disease.

**Figure 1.**
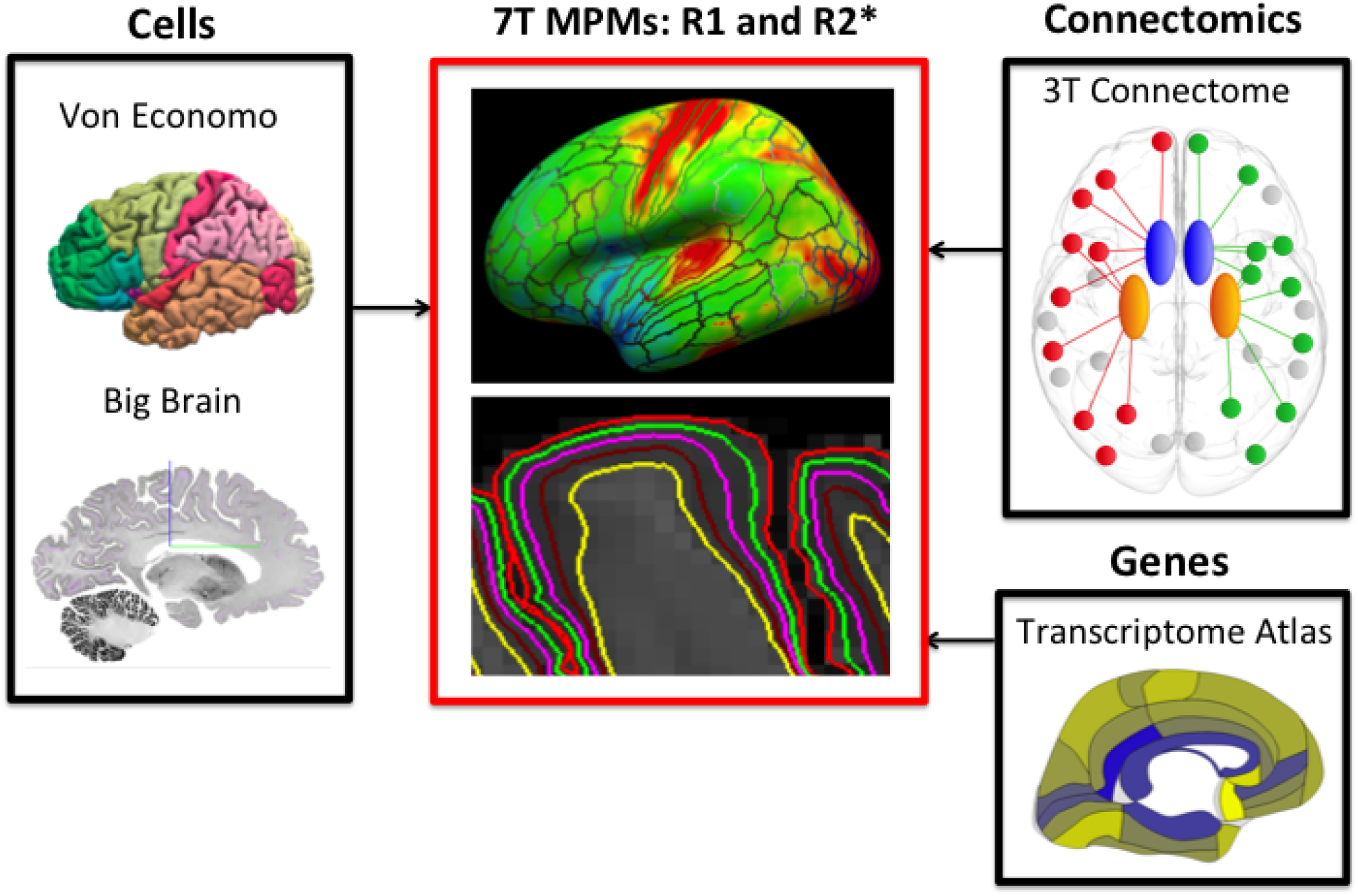
Assessment of layer-specificity of MRI. Exploring R1 and R2* quantitative 7T MRI across cortical depths using cytoarchitectonics, connectomics based on diffusion weighted imaging (DWI) with ultra-strong gradients and regional gene expression.

## Results

### R2* and R1-based myelination patterns across cortical regions and depths

The first aim of this study was to use UHF qMRI to reproduce known myelination patterns across both primary sensory and association cortices. We therefore expected that both effective transverse relaxation rate (R2*) and longitudinal relaxation rate (R1) would be higher in primary sensory areas, such as the primary visual cortex VI, than the rest of the cortex. To this aim, R2* and R1 were sampled at 50% equi-volume cortical depth and averaged across all participants. Visual inspection of R1 and R2* revealed high values in the motor and auditory cortices and in the primary visual area V1 (Fig. 2) consistent with post-mortem histology and cortical myelination patterns reported for 3T T1-weighted/T2-weighted images (Glasser et al., 2016), 7T R1 maps (Sereno et al., 2013), 3T R1 maps (Haast et al., 2016) and R2* maps (Marques et al., 2017).

**Figure 2.**
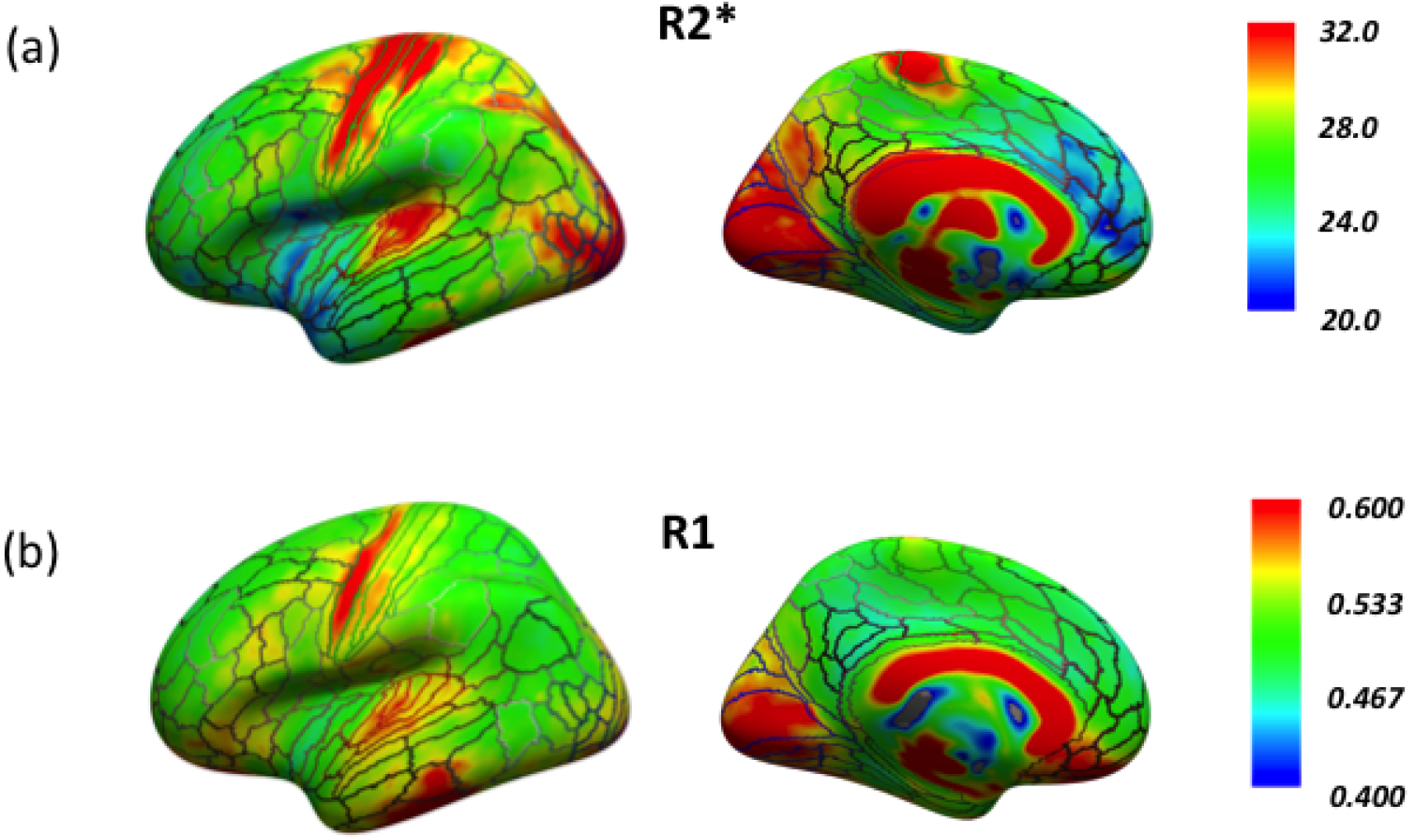
R2* and R1 values sampled at 50% cortical depth projected on Freesurfer average inflated cortical surfaces. (a) R2* (in s^-1^) (b) R1 (in s^-1^). Regions of interest (ROIs) from the human connectome project multi-modal parcellation 1.0 (HCP-MMP 1.0) atlas are outlined on the inflated surfaces.

We expected that the myelination patterns estimated at 7T for the visual, sensorimotor, auditory and parietal cortices would be comparable to those at 3T (Sereno et al., 2013; Glasser et al., 2016). This allowed us to directly compare the same HCP-MMP 1.0 atlas based ROIs at 3T and 7T.

For the visual cortex R1 was highest for primary visual cortex (V1), consistent with previous T1w/T2w ratio maps and R1 maps at 3T, however the middle temporal area (MT), which is reported as heavily myelinated at 3T showed lower R1 values (Sereno et al., 2013; Glasser et al., 2016) compared to other regions in the visual cortex. These patterns were generally consistent across depths (Fig. 3 (a)). For R2* both V1 and region MT values were greater nearest the pial surface, and then initially decreased in the middle depths before increasing between the middle and deep depths reaching highest values at the GM/WM boundary (Fig. 3 (b)).

**Figure 3.**
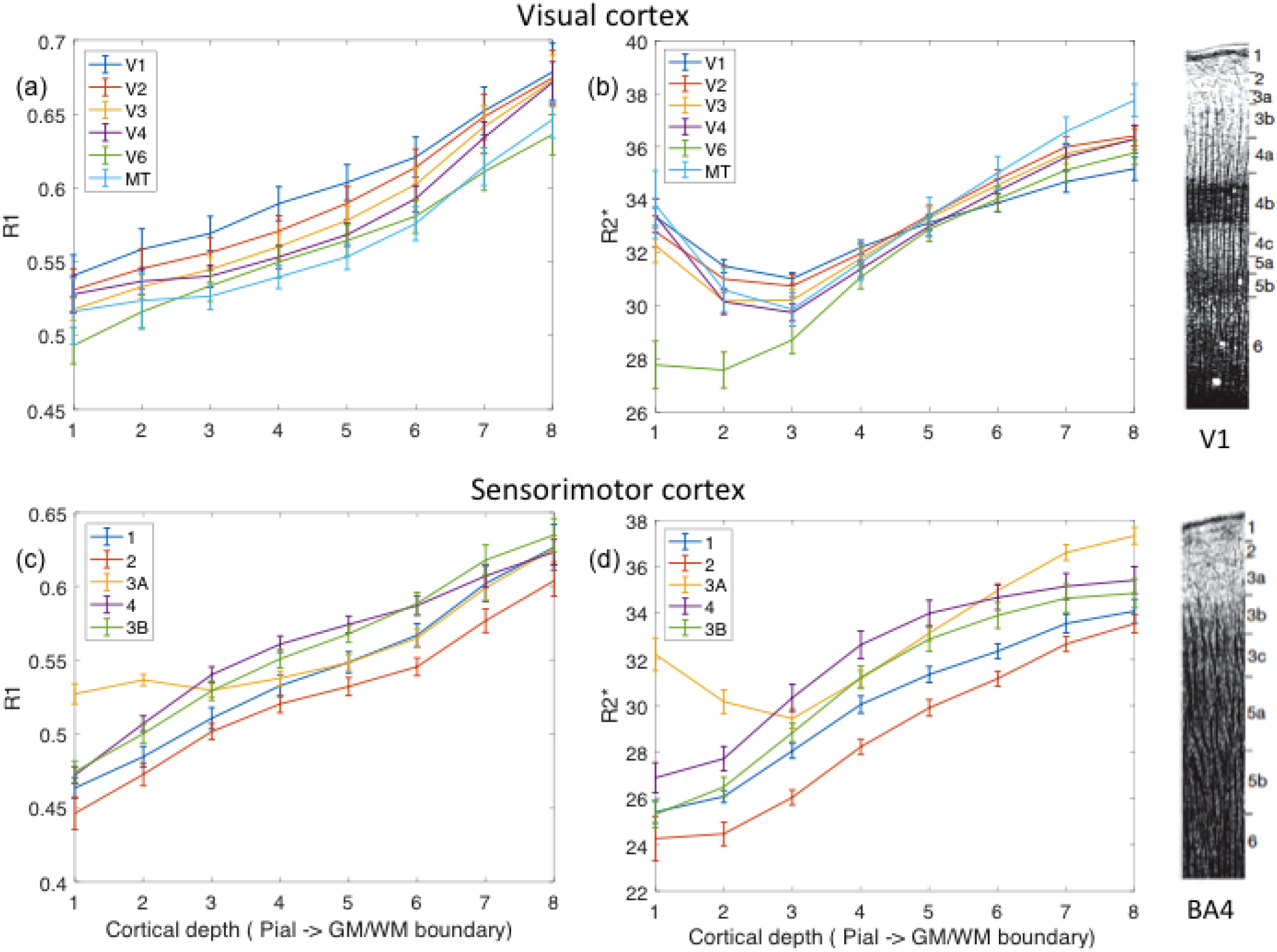
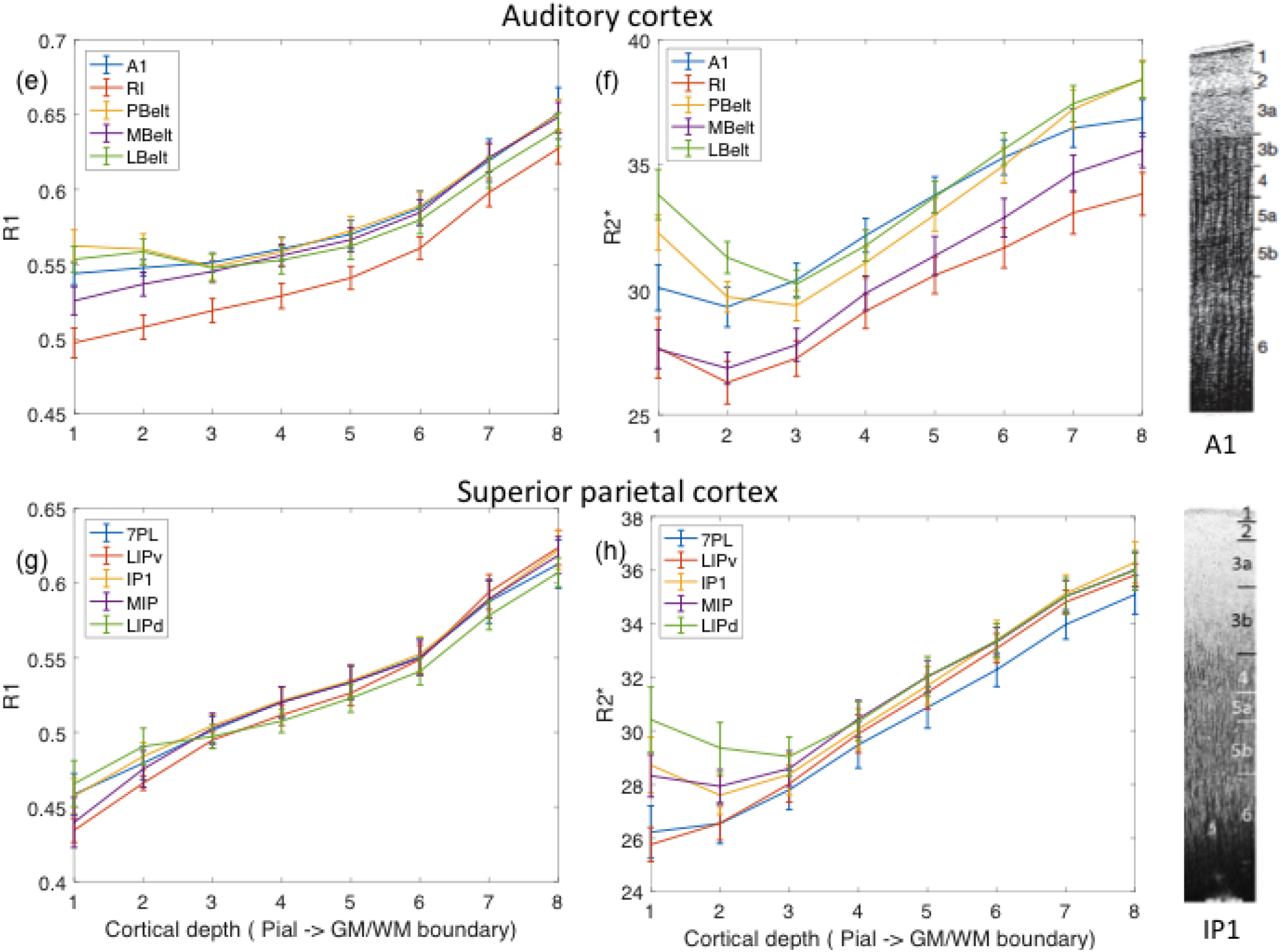
R1 and R2* profiles across primary sensory, primary motor and association cortices. Cortical depth profile, were y-axis is MRI contrast (R1 or R2*) and x-axis is equi-volume cortical depth (1 – nearest pial surface, 8 – nearest grey matter/white matter (GM/WM) boundary), for visual cortex (a) R1 and (b) R2*, sensorimotor motor cortex (c) R1 and (d) R2*, auditory cortex (e) R1 and (f) R2* and superior parietal cortex (g) R1 and (h) R2*. GM – grey matter, WM – white matter, V1 – primary visual area, V2-V6 – visual areas 2 to 6, MT – middle temporal area. 1 – area 1, 2 – area 2, 3a – area 3a, 3b – area 3b, 4 - area 4 (primary motor cortex). A1 – primary auditory cortex, RI – retroinsular cortex, MBelt – medial belt, LBelt – lateral belt, PBelt – posterior belt. 7PL – lateral area 7P, LIPv – area lateral intra-parietal ventral, IP1 – intra-parietal 1, MIP-medial intra-parietal area, LIPd – area lateral intra-parietal dorsal. Cortical labels refer to (Glasser et al., 2016). Myeloarchitectonic profiles reproduced from (Vogt, 1919; Zilles et al., 2015) are provided for areas V1 (singulostriate – absence of inner Baillarger stripe), BA4 (astriate – Baillarger stripes cannot be delineated), A1 (unitostriate – both Baillarger stripes appear to be fused to a broad band) and IPL (bistriate – both Baillarger stripes are clearly detectable).

Sensorimotor myelination patterns were consistent with patterns of T1w/T2w ratio maps at 3T and with post mortem histology (Hopf, 1968; Geyer et al., 1996; Glasser et al., 2016). Brodmann Area (BA) 3*b* showed higher R1 values than BA 3a, with the exception of the two most superficial depths. BA 2 and BA 1 showed low values for both R1 and R2*, across all cortical depths. The values in the primary motor cortex, BA 4, were highest relative to other regions in the middle depths for both R1 and R2*. In BA 3a, R1 and R2* values increased sharply between the middle and superficial depths. This may be related to blood vessel artefacts at the pial surface causing higher R2* values in the most superficial depths due to susceptibility and flow effects (Fig. 3 (c) and (d)), particularly as R2* is expected to be more sensitive to this than R1.

For the auditory cortex R1 values for primary auditory cortex A1, para belt, posterior belt (PBelt) and medial belt (MBelt) were similar, while R2* showed better discrimination of these regions. For both R1 and R2* retroinsular cortex (R1) showed the lowest values relative to neighbouring regions. This is consistent with T1w/T2w patterns at 3T (Glasser et al., 2016). Sharp decreases in R2* were seen between the superficial and middle depths for lateral belt (LBelt) and PBelt. Again this may be related to blood vessel artefacts at the pial surface causing higher values at the most superficial layers. With this exception, patterns were consistent with the literature across cortical depths for R2* and R1 (Fig. 3 (e) and (f)).

The superior parietal cortex was chosen as an example of a multimodal association area. Using T1w/T2w at 3T (Glasser et al., 2016) reported that MIP has less myelin than lateral intra-parietal dorsal cortex (LIPd) and lateral intra-parietal ventral cortex (LIPv), but MIP has more myelin than lateral parietal 7 area (7PL) and intra-parietal 1 (IP1) area. R2* values (Fig. 3 (h)) were most consistent with this showing higher MIP values compared with 7PL and IP1 across most cortical depths, with higher LIPd values higher than MIP in the superficial and middle depths. R1 values (Fig. 3 (g)) are less consistent as LIPd and LIPv are lower than MIP across most cortical depths.

In summary the R1 and R2* values we measured using 7T qMRI for sensory, motor and association cortices were generally consistent with known patterns of myelination, supporting their validity. Therefore, we proceeded to investigate their relationship to cytoarchitectonics, layer-specific gene expression and connectomics.

### Big Brain cell staining intensity is highly correlated with von Economo cell count

Before relating post-mortem histology measures to 7T qMRI we determined the relationship between von Economo and Big Brain measures at the region of interest (ROI) level and across cortical layers. We therefore performed correlations between Big Brain staining intensity and von Economo cell count and cell size. This was done by mapping the von Economo MRI atlas onto the Big Brain data using a surface-based registration. Big Brain cortical layers were defined using a machine learning approach, as previously described (Wagstyl et al., 2019). Von Economo total cell count showed a significant positive correlation with average staining intensity (rho = 0.44, p = 0.003) at the ROI level (Fig 4. (a)), whereas von Economo total cell size did not show significant correlation with average cell staining intensity profile (rho = 0.04, p = 0.79) at the ROI level (Fig 4. (b)). Comparison between von Economo layer cell count and Big Brain layer staining intensity revealed correlations across layers 2, 3, 4 and 6. Big Brain correlations with von Economo layers 2, 3 and 6 survived Bonferroni correction (Fig 4. (c)). No Bonferroni corrected correlations were seen for cell size across cortical layers. However, von Economo layer 5 cell size was most highly correlated with Big Brain layers 2, 3 and 6 (Fig 4. (d)), although not reaching significance. The absence of correlations between von Economo and Big Brain layer 1 is unsurprising given this layer contains very few cells (von Economo, 1925). The absence of correlation for layer 5 cell count in the context of positive correlation for layer 5 cell size for von Economo suggests the properties of layer 5 are quite different from layers 2, 3, 4 and 6. Indeed layer 5 contains large pyramidal cells (von Economo, 1925) suggesting large cells (which are fewer in number) may account for these findings.

**Figure 4.**
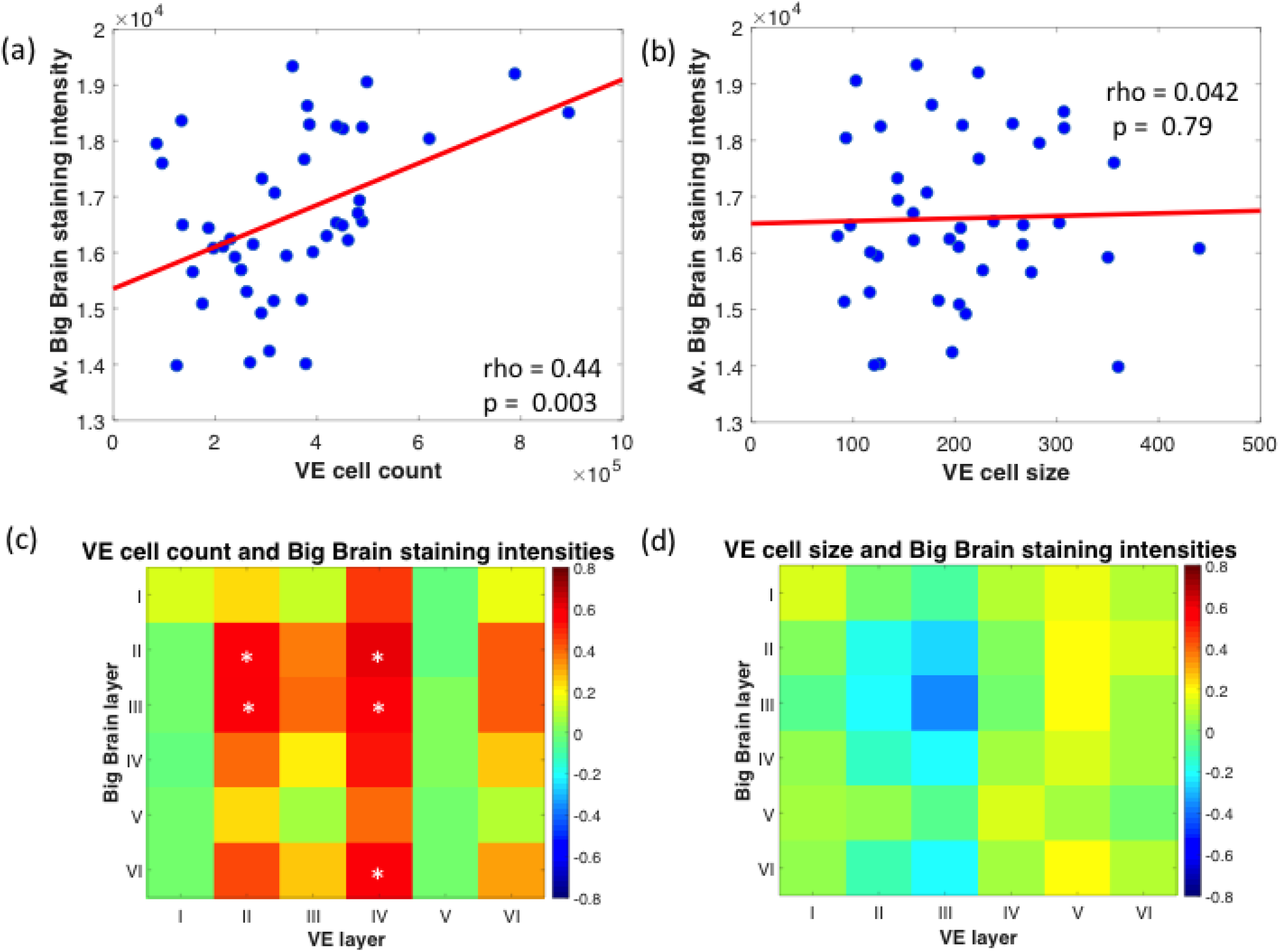
Relationship between Big Brain staining intensity and von Economo cell count and cell size. (a) Average Big Brain staining intensity plotted against von Economo (VE) cell count for each VE MRI region of interest (ROI). Each blue dot is an ROI, were the y-axis represents the average Big Brain staining intensity and the x-axis represents the average VE cell count, the red line represents a least squares linear regression line, (b) Average Big Brain staining intensity against VE cell size (μm) for each VE ROI. Each blue dot is an ROI, were the y-axis represents the average Big Brain staining intensity and the x-axis represents the average VE cell size. (c) Big Brain layer staining intensity against VE cortical layer cell count, were the y-axis represents Big Brain intensity for each cortical layer I-VI and the x-axis represents VE cell count for each cortical layer I-VI. The colours represent the correlations across VE ROIs for Big Brain intensity and VE cell count (highest – red, lowest – blue) (d) Big Brain layer staining intensity against cortical layer cell size, were the y-axis represents Big Brain intensity for each cortical layer I-VI and the x-axis represents VE cell number for each cortical layer I-VI, the colours represent the correlations across VE ROIs for Big Brain intensity and VE cell count (highest – red, lowest – blue). Asterisks indicate Bonferroni corrected significant correlations.

We assessed the intrinsic autocorrelation of histological features across cortical depth in the von Economo and Big Brain data. To this end, we determined the correlation of cell numbers between different layers within the von Economo data, which revealed cell numbers in von Economo layers 2-4 and 6 were highly cross-correlated. Big Brain staining intensity defined across cortical layers was also highly cross-correlated across all layers (see supplemental Fig. 1). These correlations were statistically significant surviving Bonferroni correction for multiple comparisons.

**Supplemental Figure 1.**
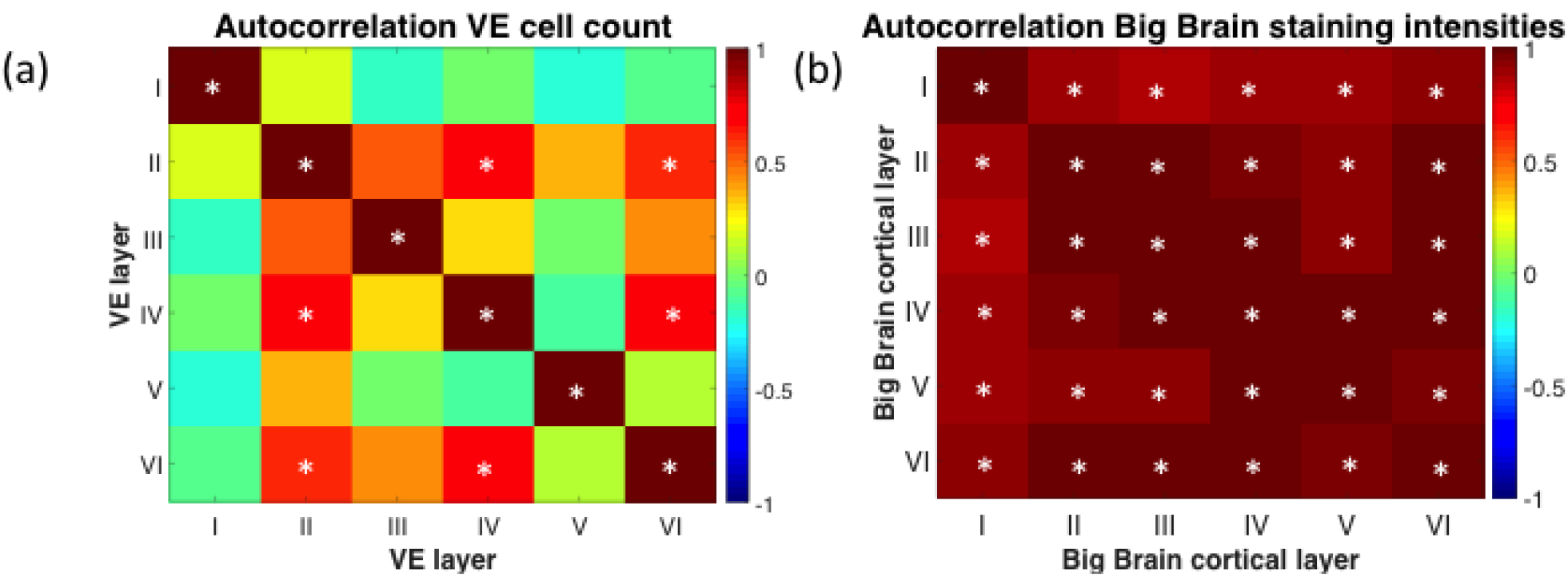
Cross-correlation for (a) von Economo (VE) layer cell count, where both the y-axis and x-axis represent cortical layer (I-VI) cell count and the colours represent correlations across VE ROIs (highest – red, lowest – blue), and (b) Big Brain cortical layer staining intensity, where both the y-axis and x-axis represent Big Brain cortical layer (I-VI) staining intensity and the colours represent correlations across VE ROIs (highest – red, lowest – blue). Asterisks indicate Bonferroni corrected significant correlations.

### R2* across cortical depths is highly correlated with von Economo cell count and Big Brain cell staining intensity across cortical layers

To examine the sensitivity of the R1 and the R2* measures to cytoarchitectonic properties, we compared R1 and R2* at the ROI level and across cortical depths with Big Brain intensity and von Economo cell count and cell size. Motivated by the relation between cyto- and myeloarchitecture and the myelin-sensitive quantitative MRI measures (Hellwig, 1993; Dinse et al., 2015) we hypothesised that UHF qMRI parameters across cortical depths would correlate with cell measures in post-mortem cortical layers, such R1 and R2* at depths near the pial surface would correlate with superficial layers, R1 and R2* at mid-depths would correlate with layer 4, while R1 and R2* at depths near the GM/WM boundary would show greater correlation with deep layers.

For the purposes of this analysis the HCP-MMP 1.0 (Glasser et al., 2016) and von Economo MRI (Scholtens et al., 2018) atlases were mapped to MPM images using surface-based registration, and eight equi-volume cortical depths were defined using Nighres (Huntenburg et al., 2018). R1 and R2* values were then sampled across ROIs and depths and averaged across participants. The Big Brain data was parcellated using the HCP-MMP 1.0 atlas and cortical layers were defined using machine learning (Wagstyl et al., 2019).

At the ROI level significant correlations were seen between R2* and von Economo cell count (rho = 0.65, p = 2.93×10^-6^) (Fig 5. (a)) and Big Brain staining intensity (rho = 0.58, p = 6.81×10^-18^) (Fig. 5 (b)). Across cortical layers Bonferroni corrected significant correlations for R2* were seen in von Economo layers 2 (across all R2* cortical depths), 3 (for mid cortical R2*, depths 3-6), 4 (across R2* cortical depths 1-7) and 6 (for R2* superficial layers, depths 2-3) (Fig. 5(c)). For Big Brain cortical layers Bonferroni corrected significant correlations were seen across all layers and all depths of R2*. In keeping with previous Big Brain and von Economo comparisons, correlations for R2* were absent for von Economo layers 1 and 5, similarly for Big Brain cortical layers correlations with R2* were lowest for layers 1 and 5. This is likely due to the small number of cells in layer 1 and the large size of pyramidal cells in layer 5 discussed previously (Fig. 5 (d)).

**Fig. 5.**
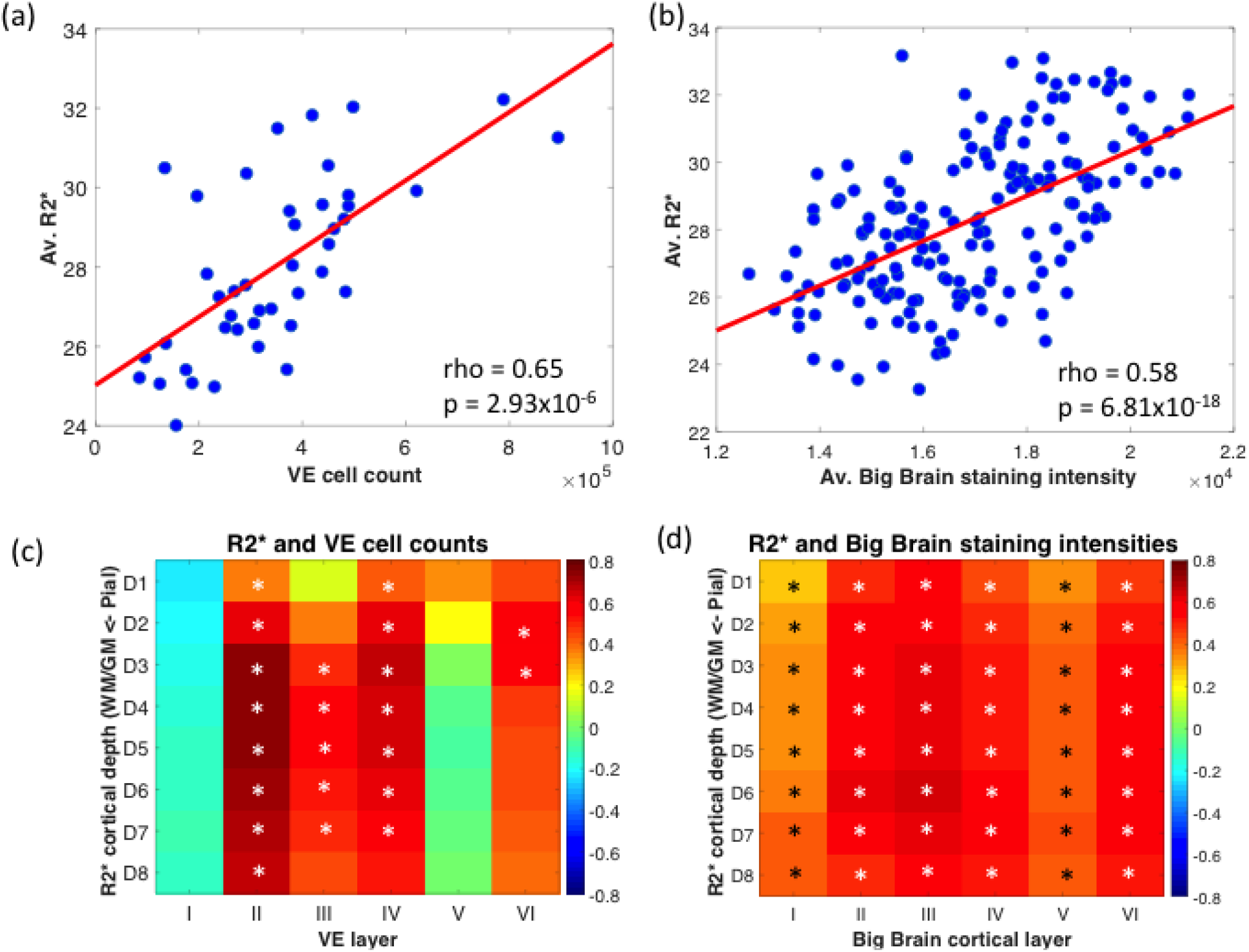
Relationship between R2*, von Economo cell count and big brain cell staining intensity. (a) Average R2* against von Economo (VE) cell count for each VE MRI region of interest (ROI). Each blue dot is an ROI, were the y-axis represents the average R2*, for each ROI across participants, and the x-axis represents the average VE cell count, the red line represents a least squares linear regression line (b) Average R2* against average Big Brain intensity for each HCP-MMP 1.0 MRI ROI. Each blue dot is an ROI, where the y-axis represents the average R2*, for each ROI across participants, and the x-axis represents the average Big Brain staining intensity. (c) R2* across cortical depths against von Economo cortical layer cell count, where the y-axis represents R2* for each equi-volume cortical depth 1-8 and the x-axis represents VE cell number for each cortical layer I-VI, the colours represent the correlations across VE ROIs for R2* and VE cell count (highest – red, lowest – blue) (d) R2* across cortical depths against Big Brain cortical layer intensity, where the y-axis represents R2* for each equi-volume cortical depth 1-8 and the x-axis represents Big Brain staining intensity for each cortical layer I-VI, the colours represent the correlations across HCP-MMP 1.0 ROIs for R2* and Big Brain staining intensity (highest – red, lowest – blue). Asterisks indicate Bonferroni corrected significant correlations.

For the R1 analysis, no Bonferroni corrected significant correlations were seen at the ROI level for von Economo cell count or Big Brain staining intensity. Across layers no Bonferroni corrected significant correlations were seen for the von Economo data (see Supplemental Fig. 2). Removal of frontal and temporal regions, which are more susceptible to artefact at 7T, due to off-resonance effects and RF transmit field reductions, had minimal effect on correlations with von Economo with none surviving Bonferroni correction. Bonferroni corrected significant correlations were seen for R1 and Big Brain layer intensity at depth 7 for layer 1 and depth 8 for layers 1-6.

Our findings show R2* but not R1 is highly correlated with cell count and cell staining intensities across cortical depths particularly for layers 2, 3, 4 and 6. The presence of high cross correlations between layers for both von Economo and Big Brain demonstrated previously may at least partly account for the lack of layer specificity of the observed correlations between the MRI parameters and histological measures, i.e. UHF qMRI parameters near the pial surface correlating with superficial layers but not deep layers.

**Supplemental Fig. 2.**
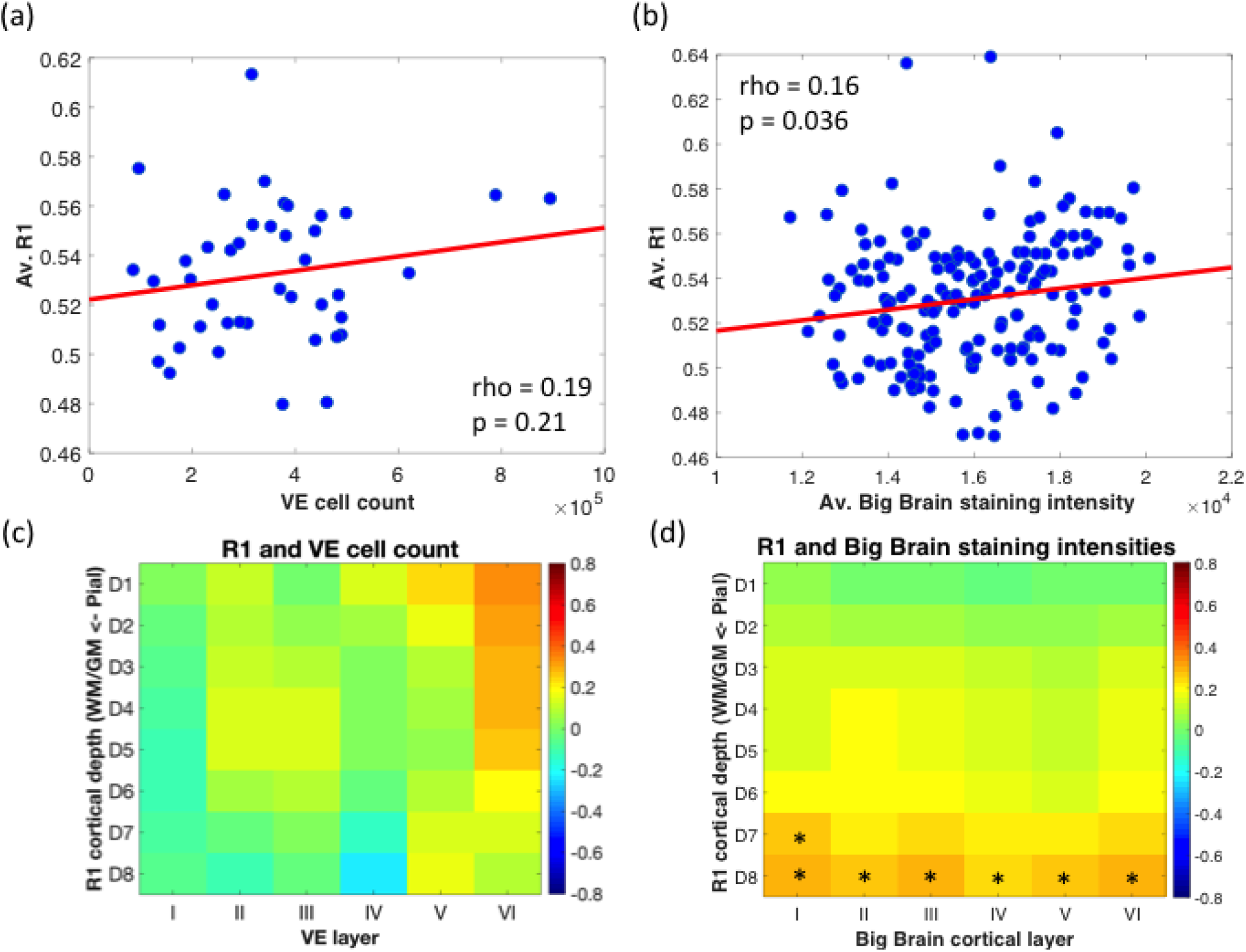
Relationship between R1, von Economo cell count and Big Brain staining intensity. (a) Average R1 against von Economo (VE) cell count for each VE MRI region of interest (ROI). Each blue dot is an ROI, where the y-axis represents the average R1, for each ROI across participants, and the x-axis represents the average VE cell count, the red line represents a least squares linear regression line (b) Average R1 against average Big Brain intensity for each HCP-MMP 1.0 MRI ROI. Each blue dot is an ROI, where the y-axis represents the average R1, for each ROI across participants, and the x-axis represents the average Big Brain staining intensity. (c) R1 across cortical depths against von Economo cortical layer cell count, where the y-axis represents R1 for each equi-volume cortical depth 1-8 and the x-axis represents VE cell number for each cortical layer I-VI, the colours represent the correlations across VE ROIs for R1 and VE cell count (highest – red, lowest – blue). (d) R1 across cortical depths against Big Brain cortical layer intensity, where the y-axis represents R1 for each equi-volume cortical depth 1-8 and the x-axis represents Big Brain staining intensity for each cortical layer I-VI, the colours represent the correlations across HCP-MMP 1.0 ROIs for R1 and Big Brain staining intensity (highest – red, lowest – blue). Asterisks indicate Bonferroni corrected significant correlations.

### R2* is highly correlated with layer-specific genes and genes related to Huntington’s disease

To further examine the relationship between qMRI parameters and layer-specific properties, we next investigated the relationship between R1 and R2* and layer-specific genes. In a similar vein to the cytoarchitectural features, we hypothesised qMRI parameters in depths near the pial surface would correlate with genes specific to layers 2 and 3, mid-depths would correlate with layer 4 and depths near the GM/WM boundary would correlate with layers 5 and 6.

For this analysis lists of genes specific to cortical layers were obtained (Bernard et al., 2012) and brain region expression data was taken from the AHBA for each ROI in the left hemisphere of the HCP-MMP 1.0 atlas, as the complete data set is not available for the right hemisphere (Glasser et al., 2016; Arnatkevic lute et al., 2019). R2* values were averaged for each ROI. For the layer-specific gene lists correlations were performed using principal component analysis (PCA), using the first component, and also mean gene expression of each list. The latter was performed in order to confirm the direction of correlation. Significant Bonferroni corrected correlations were seen for R2*averaged across layers and ROI with genes specific to cortical layer 2 (rho = −0.73, p = 2.24×10^-30^), layer 3 (rho = −0.78, p = 1.32×10^-36^), layer 4 (rho = 0.76, p = 3.51×10^-34^) and layer 5 (rho = −0.80, p = 1.01×10^-39^), but not layer 6 (rho = 0.086, p = 1.26; Fig. 6a-d). Similarly significant correlations were seen for R2* across cortical depths (averaged across the entire brain) and genes specific for layers 2, 3, 4 and 5, but not for layer 6 (Fig. 6e). Significant correlations for both PCA and mean gene expression analyses were in the same direction. No correlations between ROI averaged R1 and layer-specific genes were significant (p>0.2 uncorrected for all correlations). In order to assess whether specific genes were driving the correlations with R2*, for each individual gene in the layer-specific gene lists correlations were performed between gene expression and ROI averaged R2*. This did not identify a single gene or group of genes that were driving these results (see supplemental information).

**Fig. 6.**
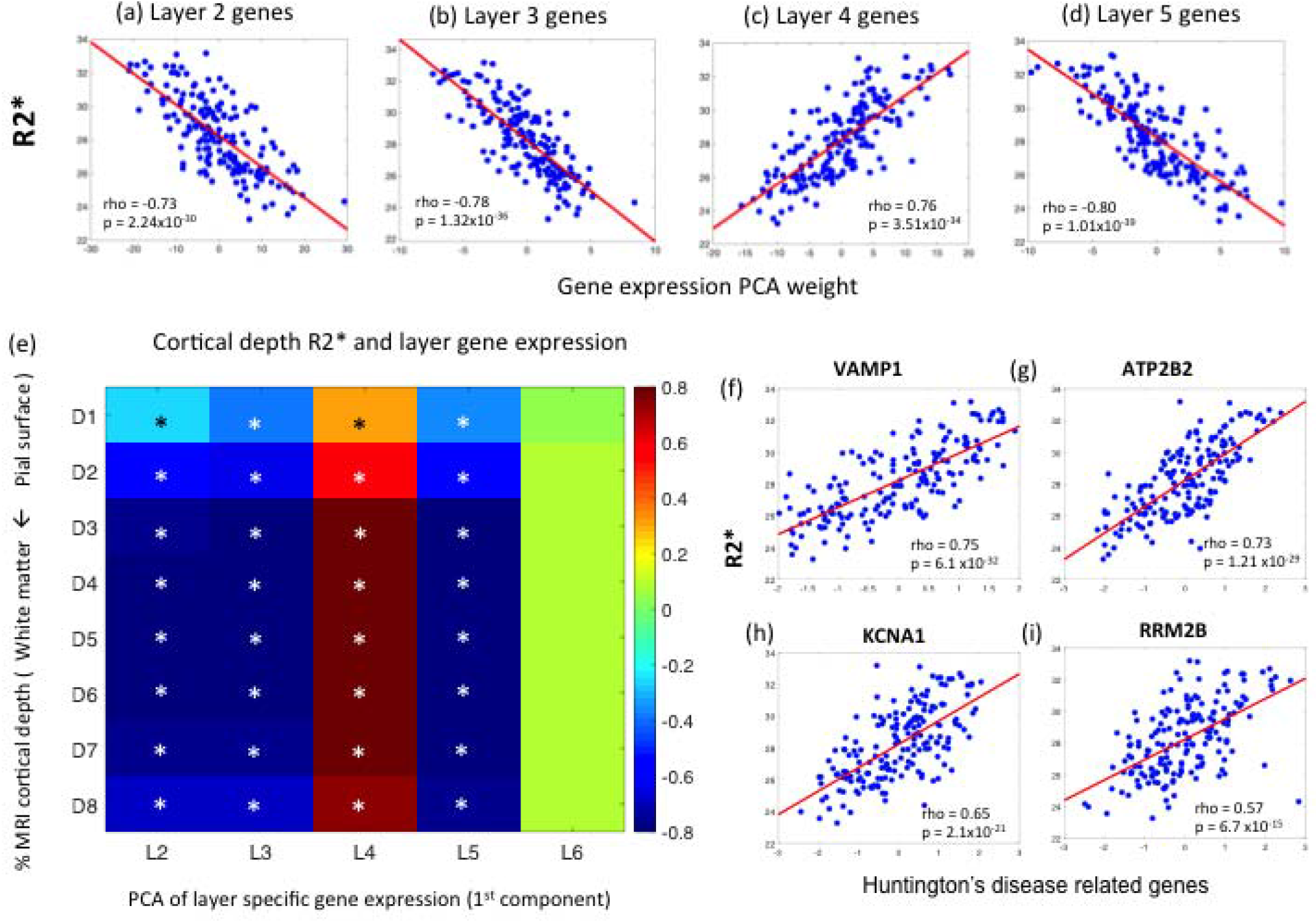
7T MRI R2* and cortical layer-specific genes and genes implicated in Huntington’s disease. R2* against (a) layer 2 genes, (b) layer 3 genes, (c) layer 4 genes, (d) layer 5 genes. Each blue dot represents an ROI from the HCP-MMP 1.0 atlas, the y-axis represents R2*, averaged across participants, and the x-axis represents the PCA weight for each ROI from the first PCA component of layer-specific gene expression, (e) R2* across cortical depths, where the y-axis represents R2* for each equivolume cortical depth 1-8 and the x-axis represents the PCA weight for each ROI from the first PCA component of layer-specific gene expression for layers II-VI, the colours represent the correlations across HCP-MMP 1.0 ROIs for R2* and layer-specific gene expression (highest – red, lowest – blue). Asterisks indicate Bonferroni corrected significant correlations. The most highly R2* correlated Huntington’s disease genes (f) VAMP1, (g) ATP2B2, (h) KCNA1, (i) RRM2B, where each blue dot represents an ROI from the HCP-MMP 1.0 atlas, the y-axis represents R2*, averaged across participants, and the x-axis represents the gene expression for each ROI for the named gene. The red line represents a least squares linear regression line.

As an example of how our analyses could be used to examine the mechanism in neurodegenerative diseases and to establish a standard reference in healthy volunteers, we explored the relationship between the regional expression of genes implicated in Huntington’s disease pathogenesis and R2* and R1. A list of 29 candidate genes were tested including 24 which show abnormal transcription in the cerebral cortex in HD (Langfelder et al., 2016), 4 genes which modify disease onset (Genetic Modifiers of Huntington’s Disease, 2015) and the normal huntingtin gene. Of those 29 tested 19 survived Bonferroni correction. The four most highly correlated candidate genes included RRM2B (rho = 0.57, p = 6.7×10^-15^), which modifies disease onset, KCNAl (rho = 0.65, p = 2.1×10^-21^), ATP2B2 (rho = 0.73, p = 1.21×10^-29^), and VAMP1 (rho = 0.75, p = 6.1×10^-32^) all of which show abnormal transcription in the cortex in HD (Fig. 6f-i). For R1 only one gene, OLFM1 (rho = 0.27, p = 0.01), survived Bonferroni correction. Results for all 29 genes tested are available in supplemental information.

### R1 and R2* across cortical depths are highly correlated with Rl/R2*-weighted white matter connections determined by DWI

The last step in our assessment examined the relationship between cortical depth dependent qMRI parameters and white matter connections estimated by DWI based tractography. We hypothesised that UHF qMRI parameters across all cortical depths would correlate with characteristics of the cortico-cortical connections, as these project across cortical layers 2-6, while qMRI parameters near the GM/WM boundary would correlate with characteristics of cortico-striatal and cortico-thalamic connections as these project to the deep cortical layers 5 and 6.

For this analysis, whole brain diffusion tractography was performed in the same participants. A white matter connectome was generated and connections were then sub-divided into cortico-striatal (C-S), cortico-thalamic (C-T) and cortico-cortical (C-C). C-C connections were further sub-divided into inter-hemispheric (inter-H) and intra-hemispheric (intra-H), as inter-H connections are more vulnerable than intra-H connections in a number of neurodegenerative diseases (Qiu et al., 2016; McColgan et al., 2017; Lanskey et al., 2018). Following a tractometry approach, streamlines were multiplied by: cross-sectional area (Smith et al., 2015) based on diffusion signal (streamline weighting); average R1 (R1 weighting); and R2* (R2* weighting). This type of tractometry is analogous to fractional anisotropy weighting performed in previous connectome studies (van den Heuvel and Sporns, 2011). Correlations were then performed for streamline-weighted connection subtypes against both R1 and R2* across cortical depths. White matter connections weighted by R1 and R2* were also correlated with R1 and R2*, respectively, across cortical depths.

The streamline-weighted connectome was not significantly correlated with either R1 or R2* across all cortical depths after Bonferroni correction (Fig. 7 (a) and (b)). However the R2*-weighted connectome showed Bonferroni corrected significance with R2* across nearly all cortical depths for cortico-striatal (depths 1-7) and intra-hemispheric (depths 1-8) white matter connections. Significant correlations were also seen for cortico-thalamic connections at superficial depths (depths 1-2). Similarly the R1 weighted connectome showed Bonferroni corrected significance with R1 across nearly all cortical depths for cortico-striatal (depths 2-8), particularly at the GM/WM boundary, and intra-hemispheric connections (depths 2-3 and 7-8) (Fig. 7 (c) and (d)).

**Fig. 7.**
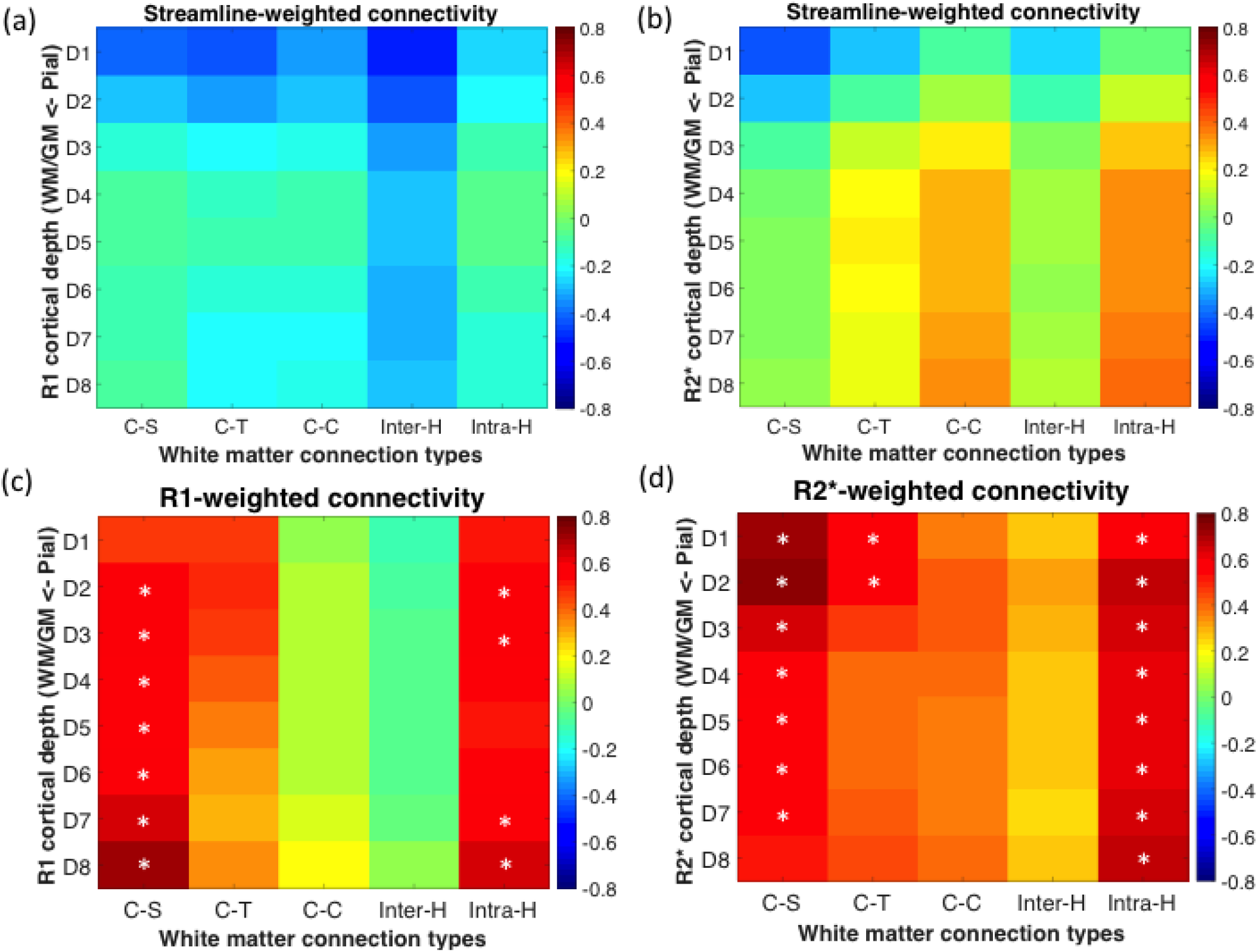
Cortical depth 7T qMRI is related to white matter connection subtypes. (a) R1 cortical depth against streamline-weighted connections, were the y-axis represents R1 across cortical depths averaged across participants and the x-axis represents streamline weighted connectivity for different white matter connection subtypes (cortical-striatal (C-S), cortical-thalamic (C-T), cortical-cortical (C-C), Inter-hemispheric (Inter-H), Intra-hemispheric (Intra-H), averaged across participants. Colours represent correlation across Killiany-Desikan atlas ROIs for R1 and streamline weighted connectivity (highest – red, lowest – blue), (b) R2* cortical depth against streamline-weighted connections, (c) R1 cortical depth against R1-weighted connections, (d) R2* cortical depth against R2*-weighted connections. Asterisks indicate Bonferroni corrected significant correlations.

## Discussion

The aim of this study was to establish the use of qMRI parameters for the study of laminar microstructure in the human brain in-vivo. For this purpose we examined the relationship between R1 and R2* measures at 7T across different cortical depths with well-established histological measures, gene-expression data and white-matter connectivity. We show that R2* across cortical depths is highly correlated with layer-specific cell number, cell staining intensity and gene expression for the von Economo, Big Brain and Allen Human Brain atlases, respectively. This highlights the sensitivity of R2* to layer-specific cytoarchitecture and gene expression. Furthermore, cortico-striatal and intra-hemispheric white matter connections weighted by R1 and R2* were highly correlated with R1 and R2* respectively across most cortical depths demonstrating the potential of linking cortical characteristics and white matter connections using 7T qMRI R1 and R2*.

### Depth profiles of myelination across cortical areas

Myelination patterns for the 7T MPMs presented here are consistent with post-mortem histology and previous 7T and 3T studies using both R1 (Sereno et al., 2013; Haast et al., 2016) R2* (Marques et al., 2017) and T1w/T2w ratios (Glasser et al., 2016) as myelin markers. Whole brain R1 maps at 700μm (Sereno et al., 2013; Haast et al., 2016) and R2* maps at 650μm (Marques et al., 2017) generated using 7T are consistent with the 500μm myelination patterns presented here with highest levels of myelin seen in the primary visual cortex, the auditory cortex and the somatosensory cortex.

When inspecting myelination profiles from the pial to the white matter surface in some cases there are distinct differences between R1 and R2*. In general R2* tends to be higher in very superficial layers, which may be caused by susceptibility effects in pial surface veins containing paramagnetic deoxygenated haemoglobin (i.e., the blood oxygen level dependent (BOLD) effect). Both the middle temporal area and V1 showed high levels of R2* consistent with high levels of myelin in these areas (Fig. 2). In contrast for R1, middle temporal area values were lower than in visual regions V2, V3, V4 and V6. Similarly in the superior parietal cortex R1 and R2* profiles diverged, such that R2* was more reflective of patterns seen in (Glasser et al., 2016). In contrast to this R1 and R2* in the somatosensory cortex were consistent and showed a distinct hierarchy across selected regions. The observed discrepancies between R1 and R2* may be due to a different contrast-to-noise ratio (CNR) of the parameter maps. The co-localisation of iron and myelin typical in cortical regions (Fukunaga et al., 2010) leads to a rather strong additive effect for the myelination contrast particularly in R2* (Stuber et al., 2014; Callaghan et al., 2015a). The increased contrast and sensitivity may make R2* a more reliable measure of myelination. For astriate (BA 4) and unitostriate (A1) areas sharp increases can be seen for R2* at different depths of the cortex in keeping with their myeloarchitectonic classification. However, bistriate regions, such as area BA 3b and the superior parietal cortex do not show the expected double peaks in their myelin profiles. This can be most likely explained by the limited spatial sampling density and consequent partial volume effects at the qMRI resolution of 500μm compared to typically smaller thickness of anatomical layers, as discussed by (Dinse et al., 2015). Challenges of precise registration, segmentation and MRI-based depth/layer definition may add further to spatial imprecisions and reduce the effective spatial resolution (Bazin et al., 2014; Trampel et al., 2019), obscuring features in myelination profiles.

At sub-millimeter resolution physiological noise and motion become more problematic. We monitored involuntary head motion by using optical prospective motion correction (Trampel et al., 2019). B1+ inhomogeneities and residual magnetisation transfer effects are also greater in UHF MRI compared to 3T. We modelled the signal dependence on relaxation parameters (R1, R2*) as mono-exponentials and neglected the orientation dependence of R2* (Rudko et al., 2014). This may have led to systematic bias and also increased inter-subject variability, since e.g. the head orientation and thus orientation dependent effects may have varied.

Image processing at UHF is also challenging. Current pipelines are typically optimised for T1-weighted images at 3T (Fischl et al., 2002) and to a much lesser extent to 7T MRI or quantitative parameter maps (Choi et al., 2019; Tabelow et al., 2019). Therefore, we had to add processing steps, such as denoising, in order to use existing pipelines. The definition of cortical layers requires highly accurate GM/WM segmentations and the use of equi-volume as opposed to equi-distant layer definition approaches (Waehnert et al., 2016), which take into account the morphology of the cortex. In order to achieve this we combined CAT12 for segmentations and Nighres software for the equi-volume layering (Huntenburg et al., 2018).

### Correlations of R1, R2*, and von Economo and Big Brain histology atlases

High correlations were observed between Big Brain cell staining intensity and von Economo cell count, but not cell size. This indicates that while cell staining intensity in the Big Brain atlas may potentially conflate cell size, cell count and cell staining inhomogeneity, cell count is the most significant contributor to cell staining, with the exception of layer 5, which shows no correlation with cell count and a modest correlation with cell size.

In comparing 7T qMRI correlations with von Economo cell count and Big Brain cell staining intensity a clear pattern emerges such that R2* across cortical regions and cortical depths shows higher correlations than R1 with post-mortem histological atlases. The relation between cytoarchitecture and the R2* and R1 parameters is believed to be largely mediated by the dependence of myeloarchitecture on cytoarchitecture (Hellwig, 1993; Dinse et al., 2015), since myeloarchitecture is known to influence macromolecular concentration and iron concentration as major MRI contrast drivers. The increased correlations of R2* over R1 with cytoarchitecture may be due to the presence of iron in multiple cell types including neurons, astrocytes and oligodendrocytes. Thus, the correlation with cell number / cell staining intensity and R2* may reflect the iron present in both oligodendrocytes and neurons (Ward et al., 2014) and macromolecules in myelin, whereas lower correlations are seen with R1 as this is mainly sensitive to macromolecules in myelin in oligodendrocytes (Stuber et al., 2014), but much less so to the iron. This is supported by a recent study that used weighted gene co-expression network analysis (WGCNA) to understand the cellular composition underlying the R2t* component of R2*. The R2t* relaxation rate constant depends solely on the cellular environment of water molecules. Using the AHBA the authors showed that R2t* was related to the regional expression of neurons and glia, including astrocytes, microglia and oligodendrocyte precursor cells (Wen et al., 2018).

A one-to-one relationship of UHF qMRI and post-mortem histology whereby UHF qMRI parameters at the pial surface would show high correlation with cells in superficial cortical layers and UHF qMRI parameters at the GM/WM boundary would show correlation with cells in the deep layers was not demonstrated. The reason for this is at least in part related to high crosscorrelations of cell counts and cell staining across cortical layers (see supplemental Fig. 1). Given the high auto-correlation across layers for both post-mortem atlases it is difficult to determine with this data alone the exact layer-specificity of qMRI. The auto- and inter-correlations between myelin, cells and UHF MRI complicate the interpretation and inferring causal factors.Auto-correlations across layers are much higher for the Big Brain than the von Economo atlas and this likely leads to greater similarity across layers with respect to R2* correlations. There are a number of potential reasons for this. Staining intensity in Big Brain differs from cell counting in von Economo in that it conflates cell size, cell density and neuropil, which can extend across layer boundaries. Big Brain layer surfaces are taken from 3-dimensional automated measurements at 20μm of a reconstructed volume with additional imperfections of section alignments. By contrast von Economo layers were manually segmented on 2 dimensional images at 1μm resolution with subsequent counting of cell bodies. An additional factor may be the ceiling effect of the cell staining in Big Brain. These factors may increase the autocorrelation of depth dependent measures in Big Brain.

### Relationship of R1, R2* and layer-specific and Huntington’s disease related genes

For cortical layer-specific genes high correlations are seen for layers 2, 3, 4 and 5, but not 6, and R2*. For layers 2, 3 and 5 correlations with R2* were negative, whereas correlation with layer 4 was positive. This is in keeping with a study at 3T examining gene expression at different cortical levels using the T1w/T2w ratio (Burt et al., 2018). In Burt et al. layer-specific gene lists were obtained from a study analysing the visual and mid-temporal cortex of post-mortem adult brains, where genes were assigned to cortical layers (Zeng et al., 2012). For our study we replicated these findings using a more extensive layer-specific gene list obtained from a study of 10 distinct cortical regions in the macaque (Bernard et al., 2012). The previously reported negative correlations between T1w/T2w ratios with layers 1-3 and positive correlations with layer 4 genes have been interpreted in the context of the thick and well defined granular layer 4 in primary sensory areas in contrast to a gradual loss of the granular layer in association cortices with progression up hierarchical levels (Burt et al., 2018). During cortical development there is a complex interplay between genes driving layer-specific development, such that genes associated with development of superficial layers suppress those involved in development of deep layers and vice versa (Greig et al., 2013). While this does not explain the R2* negative correlation with layer 2,3 and 5 specific genes and positive correlation with layer 4 genes we observed, the interplay between layer-specific genes may be a contributing factor.

As proof-of-concept of how such methods could be used to study neurodegenerative diseases, we examined the relationship between qMRI parameters at different cortical depths and the expression of genes implicated in Huntington’s disease. We demonstrate significant positive correlations of R2* and the expression of 19 genes from a list of 29 that are implicated in Huntington’s disease pathogenesis. This suggests that R2* could be used to study layer-specific pathophysiology in Huntington’s disease. Lack of correlation between layer-specific genes and R1 may again reflect the more ubiquitous nature of iron across different cell types (Ward et al., 2014) and the additive effect of iron and myelin driven contrast in R2* (Stuber et al., 2014; Callaghan et al., 2015a).

### Linking cortical grey matter and white matter

To link cortical grey matter and white matter we tested correlations between specific (R1 and R2* weighted) white matter connections with R1 and R2* across cortical depths. Significant correlations were seen for both R1 and R2* with cortico-striatal and intra-hemispheric white matter connections. For R1 the strongest correlations for cortico-striatal connections were seen at the lowest cortical depths, consistent with pyramidal tract neurons in layer 5 forming ipsilateral connections with the striatum (Shepherd, 2013), while intra-hemispheric connections showed correlations across all cortical depths for R2*, consistent with intratelencephalic pyramidal neurons present in layers 2-6 forming cortico-cortical connections (Molyneaux et al., 2007). Correlations between R2* and cortico-striatal connections were highest at superficial depths, however this may be related to the directional dependence of R2* in white matter due to microstructure orientation and myelin concentration (Rudko et al., 2014). No significant correlations were seen for streamline weighted connections, this suggests that there is a stronger relationship with R1/R2* in white matter and cortical grey matter than inter-modal comparisons using diffusion MRI based streamline weighting in white matter and qMRI (R1/R2*) measures in grey matter.

A previous study has shown that Big Brain cell staining intensities across cortical depths are strongly correlated with intra-hemispheric white matter connections between cortical regions (Wei et al., 2019). This is in keeping with our results showing high correlations between intra-hemispheric connections and R1 and R2* across cortical depths.

### A high anatomical precision framework for neurodegenerative disease

In summary, we have shown that whole brain R1 and R2* patterns acquired at 7T with 500μm resolution are consistent with myelination patterns seen at lower MRI resolutions and known post-mortem myeloarchitectonics. R2* at different cortical depths strongly correlates with layer-specific cell count, cell staining intensity and layer-specific genes. Furthermore, both cortical grey matter R2* and R1 have strong correlations with cortical-striatal and intra-hemispheric R1 and R2*-weighted white matter connections.

The qMRI and connectivity measures can provide a high anatomical precision framework identifying the origin and tracking the spread of neurodegeneration. Taking HD as an example, this is a monogenic dominantly inherited neurodegenerative disease, which causes striatal atrophy (Tabrizi et al., 2009) and loss of cortico-striatal white matter connections (McColgan et al., 2015; McColgan et al., 2017) prior to symptom onset, with subsequent cell death in cortical layers 3, 5 and 6 during the end stages of disease having been shown post-mortem (Rub et al., 2016). This paper establishes the framework to non-invasively test the following hypothesis: deep cortical layers will be affected first, in keeping with the early loss of cortico-striatal connections, followed by superficial layers and that the inter-regional patterns of cortical degeneration will be associated with the regional expression of HD related genes. Using the proposed framework cortico-striatal white matter loss can be characterised using R1-weighted white matter and linked to cortical myelin (R1) and cellular dysfunction (R2*) across cortical depths. This information can then be incorporated with the regional expression of genes involved in HD pathogenesis, which we have shown are highly correlated with R2*. In this manner we can combine measures of myelin and iron in white and grey matter (inferred using R1 and R2*) and measures of cell count across cortical depths (inferred using R2*) with pathogenic gene expression to form a comprehensive picture of the mechanism of neurodegeneration (see summary Fig. 8).

**Fig. 8.**
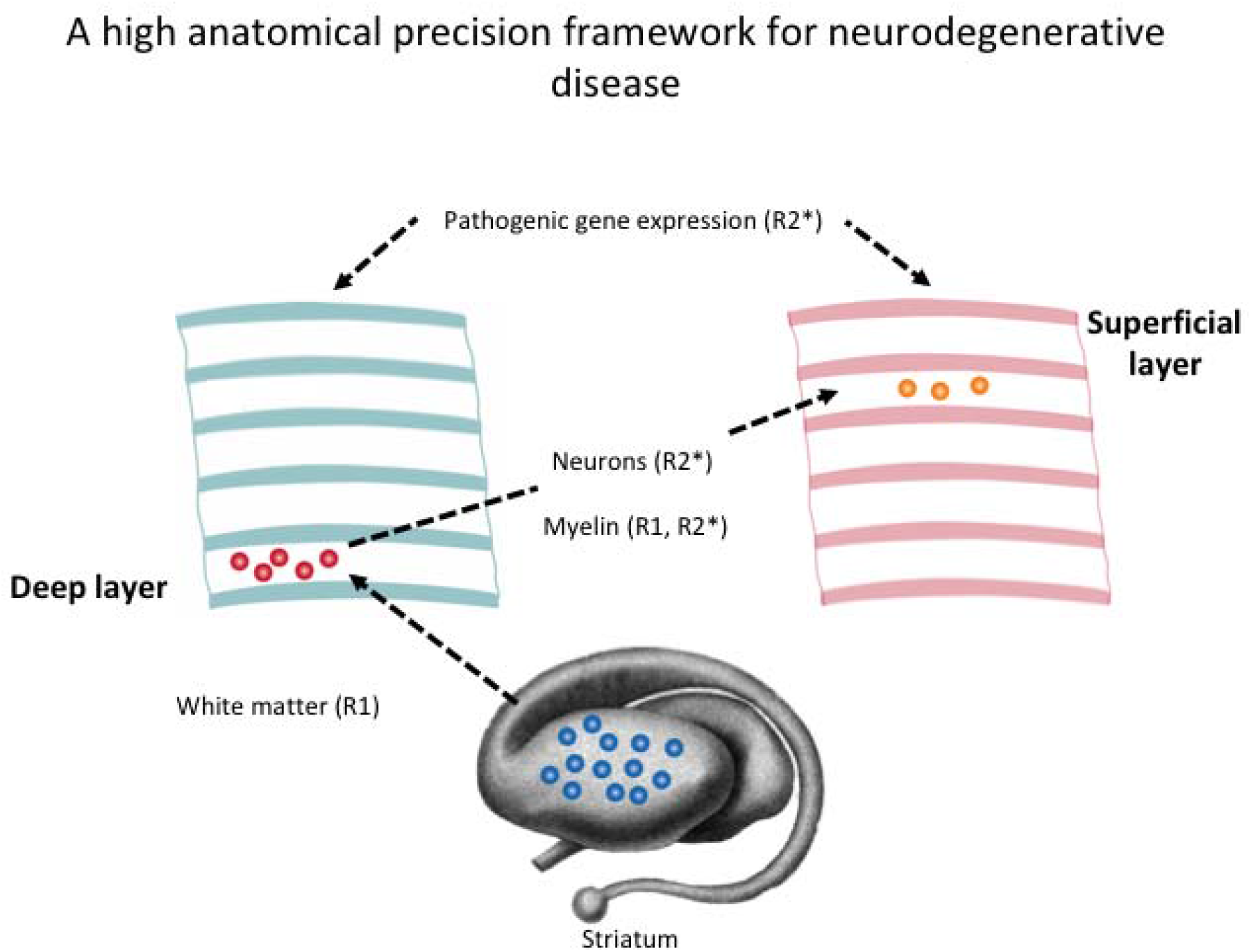
A high anatomical precision framework for neurodegenerative disease. Schematic showing how we can use the qMRI and the relationships identified in this study to combine information on white matter (R1), neuronal count at different cortical depths (R2*), myelination at different cortical depths (R1, R2*) and pathogenic gene expression (R2*) to provide a comprehensive picture of neurodegeneration.

## Materials and Methods

### Data Acquisition

Data from 10 healthy volunteers (6 females, 4 males, mean age 28±3.6 years) were acquired on a 7T whole-body MRI system (Magnetom 7T, Siemens Healthineers, Erlangen, Germany) equipped with a 1-channel transmit/32-channel radio-frequency (RF) receive head coil (Nova Medical, Wilmington, MA, USA). The MPM protocol consisted of two multi-echo fast low angle shot (FLASH) scans with T1-and PD-weighting (T1w, PDw), plus maps of the radio frequency (RF) transmit field B1^+^ and static magnetic field B0. The MPM acquisition was adapted for whole-brain coverage at 500μm isotropic resolution from methods described previously (Weiskopf et al., 2011; Weiskopf et al., 2013; Lutti et al., 2014; Trampel et al., 2019)

The PD-weighted and T1-weighted multi-echo FLASH scans were acquired with flip angles of 5° and 24° respectively, readouts of alternating polarity to give 6 echoes evenly spaced between 2.8 and 16 ms, and a TR of 25 ms for a total imaging time of 18 minutes per volume. Additional parameters were as follows: matrix size (read x phase x partition) 496 x 434 x 352, sagittal orientation, generalised autocalibrating partial parallel acquisition (GRAPPA) (Griswold et al., 2002) with acceleration factor 2 in both phase and partition directions (inner phase encoding loop), non-selective excitation with a sine-shaped RF pulse, readout bandwidth 420 Hz/pixel. The transmit voltage was calibrated by an initial low-resolution transmit field map to be optimal over the occipital lobe. Motion was monitored and corrected prospectively by an optical tracking system (Kineticor, Honolulu, HI, USA) (Callaghan et al., 2015b). For the purposes of prospective motion correction of the high resolution MPM acquisitions, each volunteer was scanned while wearing a mouth guard assembly (with attached passive Moiré pattern markers) moulded to their front teeth (manufactured by the Department of Cardiology, Endodontology and Periodontology, University Medical Center Leipzig; (Papoutsi et al., 2018)). R1, R2* and PD parameter maps were computed using the hMRI toolbox (Tabelow et al., 2019).

Diffusion weighted images (DWI) were acquired for ten healthy participants (6 females, 4 males, mean age 28±3.6 years) on a 3T Connectom (Siemens Healthineers, Erlangen, Germany) scanner (300 mT/m maximum gradient strength) using a 32-channel RF head coil for reception and a body RF coil for transmission. DWI were acquired with high isotropic spatial resolution (voxel size= 1.3×1.3×1.3 mm^3^) and four interleaved diffusion weighting shells (b=500 s/mm^2^, b=1000 s/mm^2^, b=2000 s/mm^2^, b=3000 s/mm^2^) to enable simultaneous partial volume effect (PVE) and crossing fibre modelling of the underlying voxelwise fibre populations (Jeurissen et al., 2014). DWI were acquired along 24, 36, 60 and 60 non-collinear diffusion encoding directions in the order of lowest to highest b-value shell. The diffusion encoding was achieved using monopolar diffusion weighting gradients and the diffusion weighting directions were distributed on the whole-sphere for optimum balancing and eddy current (EC)-induced distortion correction (Andersson and Sotiropoulos, 2016). A total of twenty-four non-DWI (b=0 s/mm^2^) were also acquired as baseline signal at every ten DWI (b≠0 s/mm^2^) intervals. A Centre for Magnetic Resonance Research (CMRR) single-shot, two-dimensional, multi-slice spin-echo echo planar imaging (SE-EPI) sequence (flip angle 90°, TE=65.60 ms, TR=5500 ms, partial Fourier factor=⅝, in-plane acceleration factor in phase encoding (PE) direction (GRAPPA)=2 (Griswold et al., 2002), multi-band acceleration factor in slice direction = 2 (CMRR, University of Minnesota, Minneapolis, USA. https://www.cmrr.umn.edu/multiband/, (Feinberg et al., 2010; Moeller et al., 2010; Setsompop et al., 2012; Xu et al., 2013)) with leak-block kernel reconstruction, readout bandwidth=1234 Hz/Px, effective PE bandwidth=13.256 Hz/Px, echo spacing=0.93 ms, acquisition matrix=210×212.5 (101.2% FOV phase), reconstructed matrix=162×164, number of axial slices=90, distance factor=0% and frequency selective fat suppression with both saturation RF pulses and the principle of slice selection gradient reversal was used for the DWI acquisition (Nagy and Weiskopf, 2008). The DWI were acquired with phase encoding (PE) in the anterior-posterior (AP) direction. The DWI acquisition was repeated twice to improve the signal-to-noise ratio (SNR). Five additional non-DWI (b=0 s/mm^2^) were acquired at the beginning of the DWI sequence preceded by the acquisition of five non-DWI (b=0 s/mm^2^) with reversed PE gradient polarity, i.e. posterior-anterior (PA) instead of anterior-posterior (AP) encoding direction. The collection of AP-PA non-DWI (b=0 s/mm^2^) enabled correction of susceptibility-induced geometric distortions in the DWI (Andersson and Sotiropoulos, 2016). The total DWI acquisition time was approximately 50 minutes.

An accompanying 3D magnetisation prepared rapid gradient echo (MPRAGE) image was acquired in the same session as the diffusion MRI with the following imaging parameters: voxel size =1 × 1 × 1 mm^3^, TR/TE = 2300/2.91 ms, flip angle = 9°,parallel acceleration factor (GRAPPA) = 2, field-of-view = 256 and 240 mm, matrix size = 256 x 240 x 176, sagittal slices acquired with AP phase encoding, non-selective inversion recovery with inversion time T1 = 900 ms, fat suppression and RF spoiling.

### Cortical layer construction

To enable surface based registration of cortical atlases to individual subjects the Freesurfer recon-all pipeline was used (Fischl et al., 2004) for cortical surface reconstruction. As the pipeline is designed for standard T1w MPRAGE images the following modifications were made for the 7T MPMs. A synthetic T1w image was created using Freesurfer’s mri_synthesize from R1 and PD maps. This involved scaling R1 and PD images and removing negative values. A synthetic FLASH volume with optimal white matter/grey matter contrast was created using the FreeSurfer mri_synthesize routine with TR = 20ms, flip angle = 30 degrees, TE = 2.5ms. Inputs to the routine were scaled quantitative PD and T1 maps (1/R1 volumes, with removal of a small number of negative and very high values produced by estimation errors). SPM segment (https://www.fil.ion.ucl.ac.uk/spm) was applied to the synthetic image to create a combined grey matter (GM)/white matter (WM)/cerebrospinal fluid (CSF) brain mask with a tissue probability cut-off of 0, which was used to remove the skull from the PD image. The PD image (normalised such that the average white matter intensity is at 69% (Tabelow et al., 2019)) was then subtracted from 100%, inverting the contrast and thus making it more MPRAGE-like and at the same time providing macromolecular tissue volume fraction (MTV) measures (Mezer et al., 2013). Next, Rician denoising (http://www.cs.tut.fi/~foi/GCF-BM3D) (Maggioni et al., 2013) was applied and the resulting image was used for Freesurfer cortical reconstruction. Freesurfer was then used to perform surface based registration of the von Economo (Scholtens et al., 2018), HCP-MMP 1.0 (Glasser et al., 2016) and Desikan-Killiany (Desikan et al., 2006) atlases from template to subject space.

In order to create cortical layers, first CAT12 (http://www.neuro.uni-jena.de/cat) was used to create GM and WM tissue probability maps (TPMs) from the synthetic T1w image. In CAT12 the synthetic image was spatially normalised using an affine and a non-linear registration, bias field correction was applied and the image was segmented into GM, WM and CSF (Farokhian et al., 2017). CAT12 was used instead of SPM segment as it resulted in more accurate WM segmentations for the 7T images with much higher WM tissue probabilities. In order to construct conservative GM and WM masks, the GM and WM TPMs were thresholded such that values below 1 converted to 0. This was done in order to avoid erroneous labelling of GM as WM at the GM/WM boundary. The GM TPM was manually corrected to improve minor segmentation errors. Nighres (Huntenburg et al., 2018) was used to create level set images from the GM and WM masks. The resulting outer and inner level set images were used to create 8 equi-volume layers. R1 and R2* were then sampled at each layer creating profile-sampled images. These were masked using regions of interest (ROIs) from the von Economo (Scholtens et al., 2018), HCP-MMP 1.0 (Glasser et al., 2016) and Desikan-Killiany (Desikan et al., 2006) atlases and mean R1 and R2* values were extracted for each ROI for each layer per participant. Values were then averaged across participants and only the left hemisphere was used in order to match the data from the AHBA, since it contains data from 6 left hemispheres and only 2 right hemispheres. Thus, this analysis was in keeping with previous analyses using only the left hemisphere (Arnatkevic lute et al., 2019). This resulted in matrices of 8×43 for von Economo and 8×180 for HCP-MMP 1.0 atlases.

## Diffusion MRI processing

A white matter connectome was created for each participant using anatomically constrained tractography (Smith et al., 2012) implemented in MRtrix (Tournier et al., 2012). Raw diffusion images were first visually quality controlled. Denoising (Veraart et al., 2016) and Gibbs ringing artefact removal was performed (Kellner et al., 2016) using MRtrix. FSL Eddy and Top-up were used to correct for eddy currents, susceptibility-related distortion and subject movement (Andersson and Sotiropoulos, 2016). Bias field correction was then performed using the ANTS N4 algorithm (Tustison et al., 2010). Voxelwise fibre orientation distribution were calculated using multi-shell multi-tissue constrained spherical deconvolution (MSMT-CSD) (Jeurissen et al., 2014), with group averaged response functions for WM, GM and CSF. Intensity normalisation was then performed on fibre orientation distributions (FODs) and probabilistic whole brain tractography implemented to generate 10 million streamlines. Spherical deconvolution informed filtering of tractograms (SIFT2) was used to remove biases inherent in tractography were longer connections are over-determined, streamlines follow the straightest path and lack an associated volume (Smith et al., 2013). Freesurfer was used (Fischl et al., 2004) to segment and parcellate the whole brain 3T MPRAGE image. The resulting Desikan-Killiany (Desikan et al., 2006) parcellation was used to construct the WM connection matrix. R1 and R2* weighted connectomes were also created by taking the average R1/R2* value across streamlines connecting a pair of ROIs, where higher R1/R2* were used as indicators for a higher myelination and stronger connectivity. This is analogous to fractional anisotropy weighting of streamlines, which is commonly used in white matter connectome studies, but it more directly targets the myelination of connections (van den Heuvel and Sporns, 2011). Cortico-striatal, cortico-thalamic, cortico-cortical, inter-hemispheric and intra-hemispheric connections were then extracted for each Desikan-Killiany ROI per subject and averaged across the group.

## Cell histology and genetic atlases

Cell count and cell size data across cortical layers for the von Economo atlas were taken from (von Economo, 2009), which provides cell count and cell size data for each cortical layer 1-6 in every von Economo atlas ROI. Big Brain cortical layers were defined using a machine learning approach, as previously described (Wagstyl et al., 2019). Gene expression data for the AHBA was extracted for 180 left-hemisphere regions of the HCP-MMP 1.0 atlas as detailed by (Arnatkevic lute et al., 2019). Data was available from 6 neurotypical human brains (6 left-hemisphere and 2 right hemisphere). Only data was used for the left-hemisphere as the dataset for the right hemisphere was incomplete. Genetic data processing involved six steps; gene information re-annotation, data filtering, probe selection, sample assignment, data normalisation and gene filtering. The code to run these processing steps is available at https://github.com/BMHLab/AHBAprocessing. The processed data for 180 left-hemisphere regions of the HCP-MMP 1.0 atlas is available at https://doi.org/10.6084/m9.figshare.6852911. Lists of genes specific to cortical layers 2-6 were obtained from the supplementary information of (Bernard et al., 2012).

## Experimental design and statistical analysis

In order to compare von Economo measures, cell count and cell size with Big Brain staining, correlations were performed between von Economo measures across cortical layers and Big Brain staining intensity within cortical layers as defined by machine learning (Wagstyl et al., 2019).

For R1 and R2* pairwise Pearson correlations were performed between each cortical depth and each von Economo cortical layer across 43 regions of the MRI von Economo atlas. This resulted in 8×6 (8 depth in MRI x 6 layers in von Economo) correlations for each quantitative map. A Bonferroni-corrected p(corrected) < 0.05 threshold was applied to correct for multiple comparisons (implemented by a p(uncorrected) < 0.05/(8×6)). For Big Brain data cortical layer staining intensities were used, as described above. Pearson correlations were then performed between each MRI cortical depth, for R1 and R2*, and each Big Brain staining intensity value, across 180 regions of the HCP-MMP 1.0 atlas. This resulted in 8×6 correlations. A Bonferroni-corrected p(corrected) <0.05 threshold was applied to correct for multiple comparisons (implemented by a p(uncorrected) < 0.05/(8×6)). For both von Economo and Big Brain cross correlations within each atlas across cortical depths were explored in order to aid interpretation of layer-specificity and spill over effects/covariation between layers.

For the genetic analysis R1 and R2* were each averaged for each layer within each HCP-MMP 1.0 ROI. A principal component analysis (PCA) was performed on gene expression data for layers 2, 3, 4, 5 and 6 separately. For each PCA the first principle component was selected, and Pearson correlations were performed between R1/R2* per layer and the first PCA for each ROI (Bonferroni-corrected p<0.05/5). Correlations were also performed based on mean gene expression for each gene list in order to confirm the direction of correlation from the PCA analysis. Correlations were performed between R2* across cortical depth and layer-specific genes creating a matrix of 8 depths x 5 layers (for each gene set) (Bonferroni-corrected p<0.05/(8×5)). In addition, the correlation of depth-dependent R1 and R2* with 29 genes implicated in Huntington’s disease pathogenesis was studied. Genes included four age-of-onset modifiers from a Genome Wide Association Study (GWAS) (Genetic Modifiers of Huntington’s Disease, 2015), 24 from a consensus list of genes showing abnormal transcription in Huntington’s disease (Langfelder et al., 2016) and the normal huntingtin gene. The regional expression of these genes were correlated against R1 and R2* for each ROI of the HCP-MMP 1.0 atlas (Bonferroni-corrected p<0.05/29).

Forthe connectome analysis white matter connections were split into 5 groups (cortico-striatal, cortico-thalamic, cortico-cortical, inter-hemispheric or intra-hemispheric). Correlations were then performed between these white matter subtypes and MRI cortical depth across 34 Desikan regions resulting in an 8 depths x 5 connection sub-type correlation matrix and Bonferroni correction was applied for multiple comparisons (Bonferroni-corrected p < 0.05/(8×5)). This was done for a connectome weighted by streamlines, R1 and R2*.

## Acknowledgements

We would like to thank Dr Evgeniya Kirilina for her advice regarding study design. We would also like to thank the University of Minnesota Center for Magnetic Resonance Research for the provision of the multiband EPI sequence software. We would like to thank C. Rüger and R. Haak (Department of Cardiology, Endodontology and Periodontology, University Medical Center Leipzig) for manufacturing the mouth guards for the optical prospective motion correction.

## Conflicts of Interest

The Max Planck Institute for Human Cognitive and Brain Sciences and Wellcome Centre for Human Neuroimaging have institutional research agreements with Siemens Healthcare.

NW was a speaker at an event organized by Siemens Healthcare and was reimbursed for the travel expenses.

## Funding

The research leading to these results has received funding from the European Research Council under the European Union’s Seventh Framework Programme (FP7/2007-2013)/ERC grant agreement n° 616905. NW received funding from the BMBF (01EWI711A & B) in the framework of ERA-NET NEURON, and from the NISCI project funded by the European Union’s Horizon 2020 research and innovation programme under the grant agreement No 681094, and the Swiss State Secretariat for Education, Research and Innovation (SERI) under contract number 15.0137

